# Extensive intrachromosomal duplications in a virulence-associated fungal accessory chromosome

**DOI:** 10.1101/2024.09.16.611982

**Authors:** Jelmer Dijkstra, Anouk C. van Westerhoven, Lucía Gómez-Gil, Carolina Aguilera-Galvez, Giuliana Nakasato-Tagami, Sebastien D. Garnier, Masaya Yamazaki, Tsutomu Arie, Takashi Kamakura, Takayuki Arazoe, Antonio Di Pietro, Michael F. Seidl, Gert H.J. Kema

## Abstract

Filamentous fungi have evolved compartmentalized genomes consisting of conserved core regions and dynamic accessory regions, which aid the adaptation to changing environments including the interaction with host organisms. In the *Fusarium oxysporum* species complex, accessory regions play an important role during infection and it has been reported that these regions undergo extensive duplications, however, it is currently unknown how such duplications shape accessory regions. Moreover, the function of accessory regions apart from encoding virulence effectors is not completely understood. Here we determined the karyotype of *F. oxysporum* Tropical Race 4 (TR4), which causes the ongoing pandemic of Fusarium wilt of banana (FWB). We show that the single accessory chromosome of TR4 isolate II5 has undergone extensive intrachromosomal duplications, resulting in triplication of the chromosome size compared to other closely related TR4 strains. By obtaining mutant strains that have lost the accessory chromosome, we demonstrate that this chromosome is dispensable for vegetative growth but is required for full virulence on banana. Lastly, we found that the loss of chromosome 12 co-occurs with structural rearrangements of core chromosomes, which are generally co-linear between members of the *F. oxysporum* species complex. Together, our results provide new insights into the chromosome dynamics of the banana infecting TR4 lineage of the *F. oxysporum* species complex.

**Significance:** *Fusarium oxysporum* is a major fungal plant pathogen that causes vascular wilt disease on a wide variety of agronomically important crops. A current epidemic of Fusarium wilt of banana (FWB), caused by tropical race 4 (TR4), poses a major threat to global banana production and threatens food security in tropical and subtropical regions where banana is an important staple crop. Controlling TR4 requires a better understanding of the molecular mechanisms underlying pathogenicity, including the evolution of pathogenicity-related accessory regions. Here we demonstrate that intrachromosomal duplications are a key mechanism of accessory chromosome evolution in the *F. oxysporum* species complex. We identified a single accessory chromosome and show that TR4 mutants that lost this accessory chromosome display significantly reduced virulence on banana plants. Our results provide insight into the evolution of accessory chromosomes in the *F. oxysporum* species complex, underscore their importance in pathogenicity, and provide new clues for the development of resistant banana plants.

## Introduction

Many filamentous pathogens have evolved compartmentalized genomes divided in conserved core regions and dynamic adaptive accessory regions^1–5^. These accessory regions likely evolved to facilitate adaptation to changing environments and host interactions^1,6^. Accessory regions show distinct sequence characteristics compared to their core counterparts, including a higher sequence variation, abundance in transposable elements (TEs), different codon usage, and lower gene density^3,7–9^. Moreover, accessory regions are usually enriched for histone modifications associated with facultative heterochromatin^10–13^. Accessory regions can either be embedded in core chromosomes or comprise entire independent chromosomes^14,15^.

Although the function of accessory regions is not always clear^16,17^, many have been found to play an important role in the interaction between fungi and their hosts^14,17–22^, often by encoding secreted effector proteins^3,23^. The compartmentalization into accessory regions or chromosomes may also facilitate the horizontal transfer of beneficial traits, as natural populations of different fungal species bear signs of such events^24–27^. Moreover, experimental horizontal chromosome transfer has been achieved in several species^14,25,26,28–30^. Despite these recent advances, many questions on the origin and evolution of accessory regions remain unanswered^9^.

Members of the *Fusarium oxysporum* species complex are well-known fungal pathogens containing diverse sets of accessory chromosomes and regions^23^. The accessory regions found in *F. oxysporum* strains display characteristics common to analogous regions in other fungi^15,23^, including the presence of effector genes that play an important role in virulence^14,17,21,28,31^. Furthermore, horizontal acquisition of accessory chromosomes in a non-pathogenic strain can enable the recipient strain to infect a new plant species^14^. Importantly, accessory effector profiles are often shared between strains infecting the same or a similar host^32,33^. Loss of accessory chromosomes has been experimentally induced in several *F. oxysporum* strains^17,21,29^. For example, the induced loss of a small accessory chromosome in *F. oxysporum* f.sp. *radicis-cucuminerum* resulted in complete loss of virulence on cucurbit hosts^29^.

*Fusarium oxysporum* strains infecting banana, the causal agents of Fusarium Wilt of Banana (FWB), are separated into three different races (race 1, race 2, and tropical race 4) based on their virulence profile against different banana cultivars^34^, and contain a diverse set of accessory regions^15^. The current FWB epidemic is caused by tropical race 4 (TR4) that can infect various banana varieties including Cavendish, which is widely used by the banana industry^35^. TR4 isolates are highly similar and have recently been reclassified into the species *F. odoratissimum*^36^. The TR4 reference strain II5 contains two large accessory regions: one is part of core chromosome 1 and the other is an independent accessory chromosome named AC12, which encodes various pathogenicity genes^15^. Previous work discovered signs of large-scale duplications in the accessory genome of *F. oxysporum* strains, including a copy number increase in the AC12 sequence in a subset of sequenced TR4 strains^15^. On the other hand, a recent independent study reported that the TR4 reference strain II5 lacks conventional accessory chromosomes and contains only accessory regions attached to core chromosomes^37^, which is in contrast to most thus far studied *F. oxysporum* isolates that can infect banana and other hosts^14,15,17,21,29^. Given the importance of TR4 for global banana production, these findings raise important questions about the karyotype of TR4 strain II5, the evolution of accessory chromosomes, and their roles in virulence. Here, we show that TR4 strains have an independent accessory chromosome named AC12. In the reference strain II5, AC12 is three times larger than previously reported^15^ and has evolved through repeated intrachromosomal duplication. We further show that mutant strains without AC12 show attenuated virulence on banana plants as well as reduced abiotic stress responses.

## Results

### Intrachromosomal duplications cause size expansion of accessory chromosome 12 in a TR4 strain

Accessory chromosomes in *F. oxysporum* are known to encode many predicted secreted effector proteins, play a role in virulence, and evolve by segmental duplications^14,15,38,39^. The accessory chromosome AC12 of the TR4 reference strain II5 encodes 290 predicted genes. These include 13 putative secreted effectors such as the Secreted In Xylem (*SIX*) genes *SIX9* and *SIX13*^40^, suggesting a putative role of AC12 in pathogenicity on banana. Here we confirmed these results and found that AC12 also contains genes with predicted functions in high osmolarity and cell wall stress response (Fig. 1a). Chromosome alignments showed a high similarity between AC12 from the TR4 strains II5 and M1 (Fig. 1b), which were assembled into approximately the same size (1.11 Mb for strain II5; 1.08 Mb for strain M1). However, we previously noted that AC12 from II5 has a two-to-four-fold increase in copy number compared to the genome-wide average, which is absent in AC12 from M1 (Fig 1b,^15^). Importantly, while AC12 is present in all 34 previously sequenced TR4 isolates^15^, in most of them no copy number polymorphism was detected (Fig. 1c), but we observed a copy number increase similar to that of II5 in four TR4 isolates from the Middle East (Fig. 1c). However, these four isolates are not closely related to II5, based on single-nucleotide polymorphisms on the core chromosomes, suggesting that the copy number increase of AC12 has occurred independently multiple times (Fig. 1c).

**Fig. 1.**
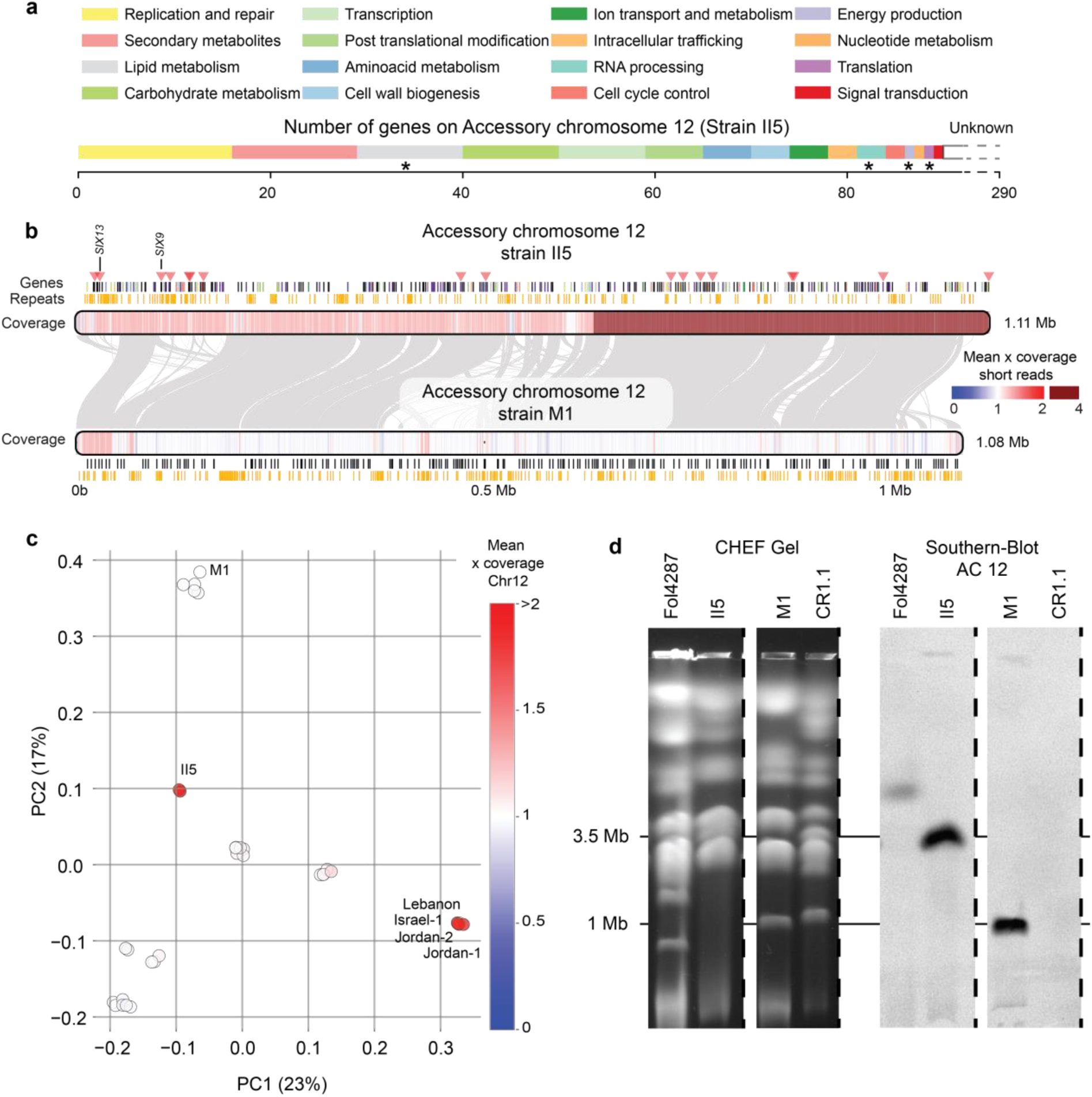
– Composition of accessory chromosome 12 (AC12) in the TR4 reference strain II5. **a**) Predicted functional categories of the genes located on AC12. Most of the genes (200) are of unknown function. Asterisks indicate gene categories significantly enriched on AC12 compared to the rest of the genome (Fisher-exact test, P < 0.05). **b**) Chromosome alignment of AC12 from isolates II5 and M1. Note that the accessory chromosomes of both strains are highly similar. Positions of predicted effector genes, including two Secreted In Xylem genes (*SIX9* and *SIX13*), are indicated by red triangles. The short-read coverage indicates a copy number increase (dark red) in II5, whereas one copy (white) is present in M1. **c**) Principal component analysis based on single-nucleotide polymorphisms (SNPs) located on the core genome of 35 TR4 strains. A copy-number increase of AC12 is found in strain II5 and in four strains from the Middle East (red dots), which are genetically different, suggesting that the event occurred independently multiple times. **d**) CHEF gel showing size-based chromosome separation of *Fusarium oxysporum* strains *Fol*4287, II5, M1, and CR1.1 and Southern blot of the separated chromosomes with a AC12 specific probe. Note that the chromosome band hybridizing to the AC12 probe is approximately 1 Mb in M1 whereas it is around 3.5 Mb in II5. Dashes indicate cropping of the gel image. The complete gel is depicted in Fig. S13.

The observed copy number increase of AC12 in II5 could be explained either by the presence of multiple separate copies of the AC12 chromosome (aneuploidy) or by large-scale internal duplications. To discriminate between these two hypotheses, we performed karyotyping of sequenced *F. oxysporum* strains using Contour-Clamped Homogeneous Electric Field (CHEF) gel electrophoresis (Fig. 1d). We included two TR4 strains, II5 showing a copy number increase, and M1 without a copy number increase (see Fig. 1b,c). We also included a banana pathogenic R1 strain CR1.1 carrying a 1.57 Mb accessory chromosome that is not homologous to AC12^15^. As a size standard, the well characterized tomato pathogenic isolate *Fol*4287^14^ was included. The CHEF gels showed the presence of a small chromosome (≤1.5 Mb) in both the M1 and CR1.1 strains, corresponding to the expected sizes of the small accessory chromosome assemblies (1.08 Mb for strain M1; 1.57 Mb for strain CR1.1; Fig.1d). By contrast, II5 lacks a chromosome band at the position of the expected size for AC12 (1.1 Mb). Southern blot analysis using a probe specific for AC12 showed a distinct hybridizing signal at the position coinciding with the smallest chromosome band of 1.08 Mb in M1, whereas in strain II5 the signal coincided with a chromosome band of a much larger size around 3.5 Mb (Fig. 1d). Importantly, the AC12-specific probe did not show a hybridization signal in R1 strain CR1.1, indicating that the signal is indeed specific for the TR4 accessory region (Fig. 1d). Thus, the size of AC12 in isolate II5 is approximately three times larger than in M1 (1.1 Mb). Together with the observed differences in chromosome coverage^15^ (Fig. 1b), these findings demonstrate that AC12 has undergone extensive intrachromosomal duplications in strain II5.

It has recently been reported that TR4 strain II5 lacks conventional accessory chromosomes^37^, suggesting that the analyzed AC12 is not present as a separate chromosome. However, we do find a separate scaffold in the published TR4 genome assembly, which has been reported to lack accessory chromosomes^37^, that is homologous to AC12 in strain II5 and M1 (scaffold 101, Fig. S1). Additionally, the CHEF gel clearly demonstrates an independent accessory chromosome in TR4, either as the smallest chromosome (M1) or as a larger chromosome that underwent intrachromosomal duplications (II5). Thus we concluded that TR4 isolates carry conventional accessory chromosomes.

The fact that the intrachromosomal duplications are collapsed in the genome assembly (1.1 Mb in the assembly vs. ∼3.5 Mb observed by CHEF analysis; Fig. 1b) indicates that these duplications are highly similar, thus preventing their separation during initial genome assembly. To unfold the assembly, we attempted to separate the duplicated reads but were unable to further improve the assembly due to the lack of sequence variation. The high similarity of the duplications strongly indicates that this evolutionary event occurred very recently. Moreover, the increase in chromosome size together with the 2x-4x increase is reminiscent of duplications caused by breakage-fusion-bridge cycles (BFB)^41^ (Fig. S2). Furthermore, the same duplications in AC12 were observed in three versions of the II5 isolate that are maintained independently for many years in three different laboratories (Fig. S3), showing that this chromosome duplication is stable.

### Accessory chromosome 12 is dispensable for vegetative growth

To investigate the function of AC12, we obtained chromosome loss mutants of TR4 strain II5. A transformant of II5 carrying a hygromycin resistance cassette inserted in the *SIX13* gene located on AC12 (II5:Δ*SIX13*) was subjected to treatment with benomyl. Seven independent colonies that had lost hygromycin resistance were obtained, and loss of AC12 was initially confirmed by PCR with AC12-specific primers (Fig. S4). The putative mutants were then sequenced to corroborate the complete loss of AC12 and check for potential additional genomic changes. Mapping of the short reads to the II5 reference genome assembly confirmed the complete loss of AC12 in all seven mutants. Importantly, we did not observe other large-scale deletions after benomyl treatment (Fig. 2a), producing similar mapping as in the wild-type strain except for AC12 loss (Fig. S5). In addition, the loss of AC12 was independently confirmed for three of the mutants by CHEF electrophoresis revealing that all three had lost the chromosome band of approximately 3.5 Mb (Fig. 2b). The hybridization signal of the 3.5 Mb band was absent in the mutants in the Southern blots, further confirming the loss of AC12 in these strains. None of the chromosome loss mutants was affected in growth on PDA plates (Fig. S6), indicating that AC12 is dispensable for vegetative growth of isolate II5. Furthermore, loss of AC12 did not impair vegetative compatibility with II5 (Fig. S7). Although AC12 loss did not affect vegetative growth or vegetative compatibility, AC12 mutants were slightly affected in their tolerance to high salinity and cell wall related stress (Fig. S8).

**Fig. 2.**
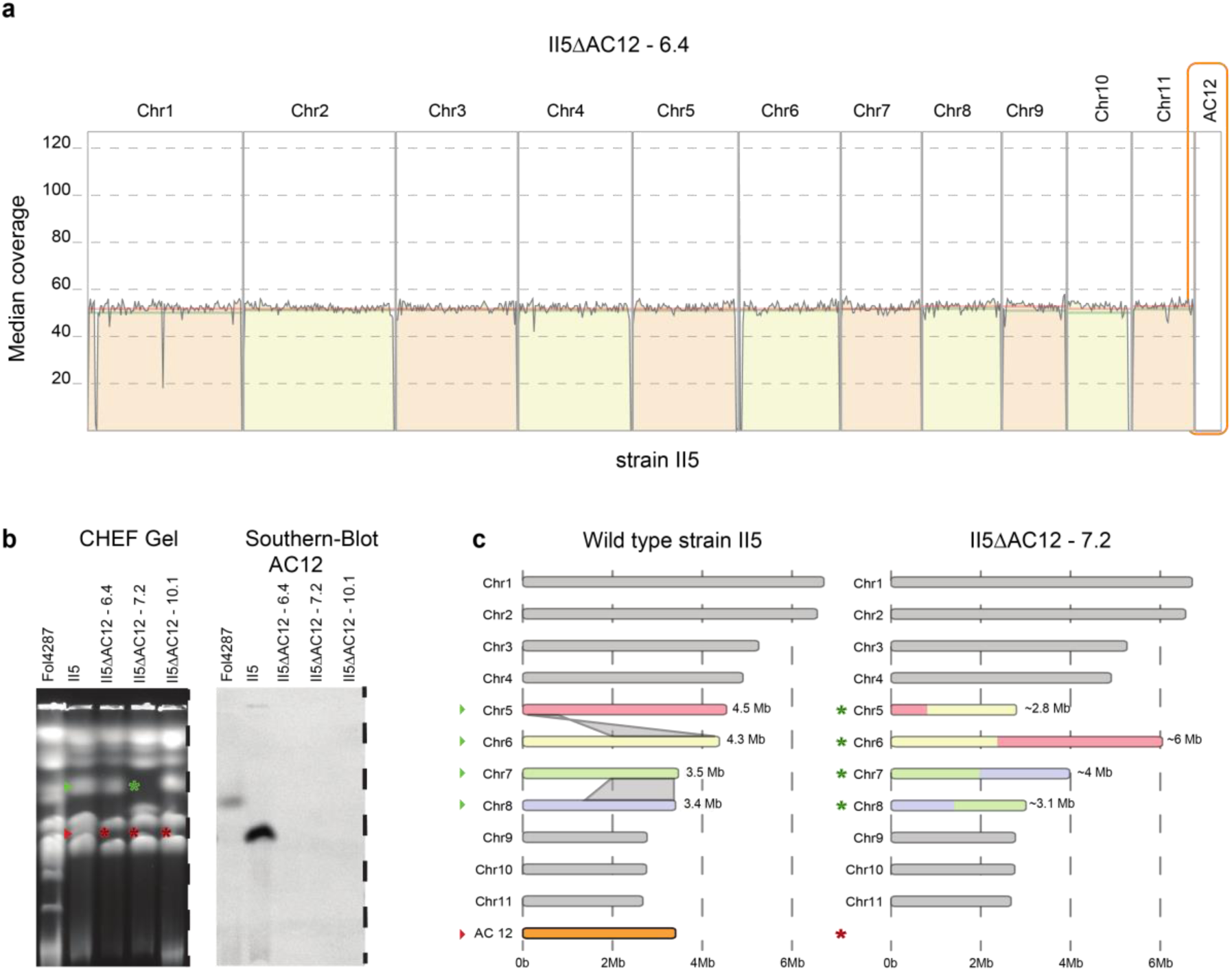
– Accessory chromosome 12 (AC12) is dispensable in strain II5. **a**) Mapping of Illumina short reads from the II5ΔAC12 mutant 6.4 to reference strain II5 demonstrates the specific loss of chromosome AC12. **b**) CHEF gel showing separation of chromosomes from *Fusarium oxysporum* strains *Fol*4287 and II5, and from three independent chromosome loss mutants II5ΔAC12 6.4, II5ΔAC12 7.2, and II5ΔAC12 10.1. Note the absence in the mutant lines of the AC12 chromosome band (left panel, red arrow and asterisks) and of the corresponding hybridization signal (right panel). Green arrow and asterisk indicate the position of a core chromosome that has undergone a rearrangement in mutant 7.2. The complete gel is depicted in Fig. S13. **c**) Schematic representation of the interchromosomal rearrangements observed in chromosome loss mutant II5ΔAC12 7.2. Translocations between chromosomes 5 and 6 and chromosomes 7 and 8 are highlighted by ribbons.

Besides the loss of AC12, the CHEF gel revealed additional karyotypic differences, most notably in mutant 7.2 which showed a change in the core chromosome banding pattern (Fig. 2b). We hypothesized that the difference observed in this mutant could be the result of interchromosomal rearrangements, since sequencing did not detect loss of genetic material other than AC12 (Fig. 2a, Fig. S5). Further analysis of short-read genome assemblies uncovered several large-scale changes. Most structural variants were either deletions (6-12 deletions per mutant) or rearrangements (4-8 rearrangements per mutant) (Fig. S9). Most notably, five of the seven mutants showed interchromosomal rearrangements between chromosomes 7 and 8. Moreover, we detected rearrangements between chromosomes 1 and 2 in mutant 7.3 (Fig. S10a), between chromosomes 5 and 6 in mutant 7.2 (Fig. 2d), and between chromosomes 6 and 7 in mutant 10.1 (Fig. S10b). The rearrangement between chromosomes 5 and 6 in mutant 7.2 was independently confirmed by PCR (Fig. S11). None of the interchromosomal rearrangements between core chromosomes caused any detectable defects during vegetative growth on PDA plates (Fig. S6). In summary, these results suggest that benomyl treatment triggers interchromosomal rearrangements between core chromosomes in addition to loss of the accessory chromosome 12 in TR4 strain II5, and that none of these changes affects vegetative growth of the fungus.

Accessory chromosomes can be horizontally transferred between different *F. oxysporum* strains through vegetative hyphal fusion^14,17,29^. However, several attempts (30 plates) to transfer AC12 carrying the hygromycin resistance cassette from strain II5 to the R1 reference strain CR1.1 or to the non-pathogenic *F. oxysporum* strain *Fo*47 failed to provide any evidence for successful chromosome transfer.

### Accessory chromosome 12 is important for virulence of TR4 on banana plants

To determine the contribution of AC12 to virulence of TR4 on banana, plants of cultivar Cavendish were inoculated with strain II5 or with three mutants lacking AC12. Interestingly, the three mutants showed a strong reduction in corm necrosis levels, a well-established indicator for virulence^42^, when compared to the wild-type strain (Fig. 3A). Although the mutants were still able to infect Cavendish, the average severity of corm necrosis was 48.9% compared to 95% in the wild type. Importantly, benomyl treatment itself did not affect the virulence on Cavendish plants (Fig. S12).

**Fig. 3.**
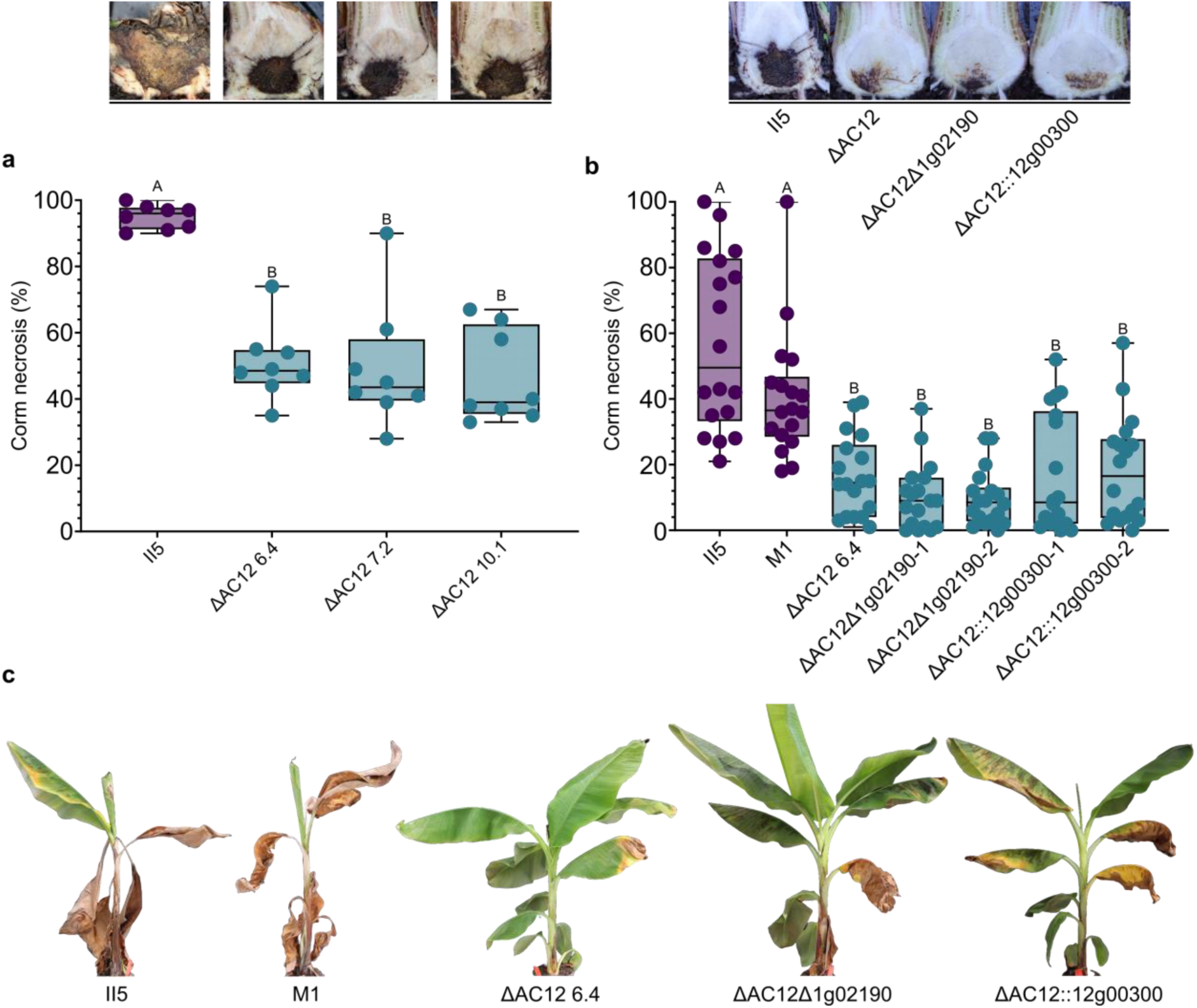
– Accessory chromosome 12 (AC12) is important for virulence of II5 on Cavendish plants. **a**) Percentage of corm necrosis of Cavendish ‘Grand Naine’ plants at 10 weeks post inoculation with reference strain II5 or with three independent AC12 loss mutants. Corm necrosis was quantified using ImageJ (n=8). Representative pictures of corms are displayed above. **b**) Percentage of corm necrosis of Cavendish ‘Grand Naine’ plants inoculated with TR4 wild type strains II5 or M1; with two independent knockout mutants in the effector gene *1g02190* obtained in the II5 ΔAC12 background (ΔAC12Δ1g02190-1 and ΔAC12Δ1g02190-2); or with two independent complemented strains where the *12g00300* gene encoding an AC12-specific effector candidate was re-introduced into an ΔAC12 background (ΔAC12::12g0300-1 and ΔAC12::12g0300-2) (n=18). Different letters indicate significant differences between treatments (Tukey-Kramer test; P<0.05). Representative pictures of corms are displayed above. **c**) Photos of external symptoms of plants analysed in panel **b**.

Individual accessory effectors were previously shown to act as virulence factors of TR4 against banana^43,44^. However, the study of some effectors has been difficult due to their duplication between the accessory region on chromosome 1 and the segmental duplications of AC12^15^. The availability of the AC12 loss mutants allows the genetic analysis of these effectors by simplifying knockout generation. To study the effect of effectors on Chr1, two independent knockout mutants of the gene *1g02190* located on the accessory region of chromosome 1, which encodes an *in planta* upregulated effector candidate, were obtained in the ΔAC12 background (ΔAC12Δ1g02190-1 and ΔAC12Δ1g02190-2). However, these effector knockout mutants did not show any additional reduction in virulence compared to the ΔAC12 mutant (Fig. 3B). By contrast, re-introducing the *12g0030*0 gene, which is unique to AC12 and encodes the only *in planta* upregulated effector, into an ΔAC12 mutant (strains ΔAC12::12g0300-1 and ΔAC12::12g0300-2), resulted in higher necrosis levels in some plants, although the difference was not significant compared with the ΔAC12 mutant. Finally, strains M1 and II5 were equally virulent, indicating that intrachromosomal duplications of AC12 do not affect virulence under greenhouse conditions. In general, external symptoms were also more severe and developed faster in the plants inoculated with the wild-type strains compared to the mutants (Fig. 3C).

## Discussion

Genome compartmentalization occurs in many filamentous fungal pathogens^2,4,5,45^. Accessory genomic regions show higher genetic variation compared to the core genome, which is thought to contribute to fast adaptation^2,4^. Effector genes important for infection of the host are often located in such dynamic accessory regions^45^. *Fusarium oxysporum* strains contain accessory regions either as independent chromosomes or as parts of core chromosomes^14,15^. These accessory regions play important roles in several *F. oxysporum* pathosystems^14,17,29^, exemplified by their ability to horizontally transfer pathogenicity on a given host plant from pathogenic to non-pathogenic isolates^14,17,28,29^.

Here, we demonstrate the presence of an independent accessory chromosome in TR4 strain II5 and we show that a recently observed difference in copy-number of AC12 in the TR4 reference strain II5^15^ is the result of remarkably large intrachromosomal duplications causing an approximate triplication of the size of this accessory chromosome. Interestingly, a similar duplication event must have occurred at least twice independently in different isolates of the clonal TR4 lineage. Since these duplications were not observed in TR4 strains isolated from Taiwan^46^, the first known origin of TR4^47^, they appear to have occurred after the initial spread of these isolates. Moreover, the high similarity of the duplicated AC12 sequences in these different isolates indicates that these events occurred recently. Due to a lack of sequence variation between these intrachromosomal duplications, they initially resulted in the assembly of a single 1.1 Mb contig^15^, instead of the 3.5 Mb chromosome detected by CHEF gel analysis. Although the initial assembly does not correctly represent the complete accessory chromosome, the assembled chromosome is highly similar to AC12 from strain M1^15^, as well as to a contig in the independently assembled UK0001 strain^48^ and to scaffold101 in an independently acquired II5 genome assembly^37^. Importantly, our results clearly demonstrate the presence of an accessory chromosome in TR4 strains, contrasting a recent report concluding that TR4 strain II5 lacks separate accessory chromosomes^37^. Similar to our findings, a karyotype analysis of three decades ago also showed the presence of a 1.1-1.2 Mb chromosome in several TR4 strains, although at that time no accompanying genome assemblies were available for the strains in this early study^46^. In addition to clarifying the karyotype of II5, we also show that AC12 is dispensable for vegetative growth on plates, further underscoring its role as accessory chromosome. Taken together, these findings firmly establish the presence of independent accessory chromosomes in TR4 strains, including strain II5, similar to those found in the majority of the previously characterized *F. oxysporum* strains.

Intrachromosomal duplications were previously reported in *F. oxysporum* after experimental horizontal transfer of accessory chromosomes^29^. It is therefore tempting to speculate that the duplications in AC12 also occurred after a horizontal transfer event. Although the mechanism driving these intrachromosomal duplications remains unknown, it could involve a process known as BFB cycles^41^. The duplications in AC12 could be explained by a different number of sequential perfect and/or complex BFB cycles, giving rise to a much larger accessory chromosome^49^. To further study the formation of the observed duplications, an accurate assembly of the full length AC12 will be essential.

The viability of mutants lacking the complete AC12 chromosome proves that it is dispensable for vegetative growth, as previously reported for accessory chromosomes in *F. oxysporum* strains infecting different host plants^14,17,29^. However, we observed a significant reduction in disease severity on banana plants after inoculations with the AC12 loss mutants, suggesting an important role of this accessory chromosome in plant infection similar to reports for accessory chromosomes in *F. oxysporum* strains infecting different plant hosts^17,21,28,31^. In addition, the growth of the AC12 loss mutants was slightly impaired under high salt and cell wall stress. Together with the putative functions of other AC12 encoded genes, the latter result suggests that this accessory chromosome likely has a wider role besides virulence. In contrast to previous observations^14,29^, the AC12 loss mutants were still able to infect the host plant and cause the typical FWB symptoms, albeit with reduced severity. This result could be explained by the presence of an additional accessory region on core chromosome 1 of isolate II5^15^, which encodes seven *SIX* homologs as well as 33 candidate effectors, suggesting an important role during infection^43,44^. Moreover, the accessory region on chromosome 1 and AC12 share many duplications, including several putative effector genes^15^. Because these duplications hamper the knockout analysis of effector candidates, the newly generated AC12 loss mutants represent a useful tool for determining the minimal set of effectors required to infect banana and facilitate the search for potential avirulence genes using resistant banana genotypes. Here we attempted to further quantify the relative contribution of AC12 and the accessory region of chromosome 1 to virulence, by generating two different effector mutants in the AC12 loss background. While knockout of an additional accessory effector gene on chromosome 1 did not further reduce virulence of the AC12 loss mutant, re-introduction into such a mutant of a single AC12-specific effector gene resulted in a slight, albeit not significant increase in virulence. Since complementation was done through ectopic integration with the native promoter, differences in gene expression might explain the partial phenotype of the complemented line. To better assess the contribution of this effector, a knockout mutant could be made in the M1 strain which lacks the duplicated copies. The contribution of AC12 to virulence of TR4 is remarkable considering the number of regions shared with the accessory region of chromosome 1^15^. Additional genes other than effectors located on AC12 might play a more prominent role in the virulence phenotype of the chromosome loss mutants than previously expected.

Besides loss of the accessory chromosome AC12, several mutant lines showed interchromosomal rearrangements of core chromosomes. These rearrangements are of interest considering the commonly observed high level of co-linearity between core chromosomes of *F. oxysporum* strains^15,38,50,51^. Although chromosomal rearrangements can have a strong impact on different functions including plant pathogenicity^52–56^, we observed no phenotypic differences in the mutants that could be associated with the core chromosome rearrangements.

Chromosome length polymorphisms have been observed for a long time in a wide array of fungi^57^, and large-scale duplications or aneuploidy have been linked to differential fitness^58,59^. Gene copy number changes are often beneficial for growth in a specific (stressful) environmental condition^59–63^. On the other hand, aneuploidy can strongly affect transcription and, in some cases, lead to proteotoxic stress^64,65^. In yeast, smaller chromosomes with fewer genes are more frequently duplicated, which might be a consequence of overexpression and proteotoxic stress^66,67^. Similarly, in yeast, aneuploidy is a driver of further genomic instability^68^. Less is known about these specific processes in filamentous fungi, despite the observed karyotype variability^54,57^. Our results showcase intrachromosomal duplication as another example of karyotype variability among field isolates of a filamentous plant pathogen. The intrachromosomal duplications appear relatively stable over time, and their independent occurrence in different isolates suggests a potential fitness benefit. It has been suggested that duplications of accessory chromosomes might be an important prelude to rapid sequence divergence and effector diversification^69^. Nevertheless, a clear fitness benefit or detriment linked to these duplications remains to be discovered.

Together, our data demonstrate the occurrence of intrachromosomal duplications of AC12 in the TR4 reference strain II5 and a contribution of AC12 to different biological processes, particularly plant infection. The observed duplications and chromosomal rearrangements indicate a high degree of genome plasticity and highlights an important mechanism of accessory chromosome diversification in fungal plant pathogens. Further knowledge on the function and dynamics of accessory regions in the TR4 lineage will be crucial for combating this devastating banana pathogen.

## Materials and methods

### Fungal growth conditions and chromosome loss induction

*Fusarium oxysporum* Tropical Race 4 (TR4) strains II5 and M1^15,70^ and newly generated II5 mutant strains were routinely cultured on PDA plates at 25 °C. Conidial suspensions of all strains were stored as glycerol stocks at –80 °C.

Chromosome loss was induced as described previously with slight modifications^17^. II5:Δ*SIX13* was cultured in M100 medium supplemented with 25 µg/mL benomyl at 150 rpm 25 °C for 4 days. Conidia were obtained by filtering the culture through two layers of sterile cheesecloth. Conidia were plated on M100 plates supplemented with 0.04% Triton X-100. Plates were overlaid with a piece of sterile filter paper and incubated at 25 °C for 2-3 days. Subsequently, the filter paper was transferred to a PDA plate with 100 µg/mL hygromycin that was incubated at 25 °C for 1 day. Afterwards, the filter paper was removed, and colonies were selected for loss of hygromycin resistance by comparison to the original M100 plate. Putative chromosome loss mutants were transferred to separate PDA plates for further validation. PCR genotyping was performed using the Phire Plant Direct PCR Master Mix (Thermo Scientific™, Waltham, Massachusetts USA). Chromosomal rearrangement between chromosomes 5 and 6 were validated using PCR as well. Primers are listed in Table S1.

### Sequencing and sequence analysis

To validate that the mutants generated by benomyl treatment lost AC12 and analyze the effect of benomyl treatment on the presence of other genomic regions, seven chromosome 12 mutants were sequenced. The mutants were grown in PDB for 4 days at 150 rpm 25 °C to obtain mycelium. Cultures were centrifugated and mycelium was freeze-dried. DNA was extracted from freeze-dried mycelium using the Masterpure™ Yeast DNA Purification Kit (LGC Biosearch™ Technologies, Hoddesdon, United Kingdom). Extracted DNA was sent for Illumina whole-genome re-sequencing by Beijing Genomics Institute. The reads were mapped against the II5 reference genome assembly^15^ using BWA mem version 0.7.17^71^. The mapping was visualized using WGS coverage plotter from jvarkit version 589510ae3^72^.

Additionally, the short reads were assembled using SPAdes version 3.13.0^73^. These assemblies were used to analyze structural variants, including chromosomal rearrangements, using mummer version 4^74^ followed by MUMandCO version 3.8^75^.

To determine the putative function of the genes located on chromosome 12, we used eggnog mapper version 2.1.12^76^. Putative effector genes were identified using effectorP version 3^77^ on the secretome determined by signalP version 5^78^. Secreted in Xylem (*SIX*) genes were identified based on homology searches of *SIX* genes from *Fol*4287 using blastP version 2.14.0^79^.

### CHEF electrophoresis and Southern blotting

Chromosomes of all strains were separated by Contour-Clamped Homogeneous Electric Field (CHEF) electrophoresis. Protoplast preparation was performed as previously described^80^, with slight modifications: 5×10^8^ fresh microconidia were germinated 14 hours at 28 °C in 200 ml of potato dextrose broth (PDB) medium, germlings were then transferred to a solution of 1.2M MgSO4 pH 5.8, containing 5% (w/v) Extralyse® enzyme preparation (Laffort, Bordeaux, France) and incubated approximately 90 minutes at 30 °C and 60 rpm. Protoplasts were filtered through a double layer of Monodur mesh (pore size 10 μm), washed with ice-cold 1 M Sorbitol and harvested after 15 minutes of centrifugation at 2,500 rpm. Subsequently, they were resuspended at a concentration of 8×10^8^/mL in STE solution (25 mM Tris-base, 50 mM EDTA, 1M Sorbitol; pH 7.5) containing 1 mg/mL proteinase K (Sigma-Aldrich, Madrid, Spain), mixed with an equal volume of 1.2% Low Melting agarose (NuSieveTM GTGTM, Lonza, Pontevedra, Spain), poured into plug casting molds (Bio-Rad, Madrid, Spain) and left to cool 15 minutes at 4 °C. These plugs were then transferred to a solution containing 1 mg/mL proteinase K in 100 mM Tris-base, 500 mM EDTA, 1% Lauryl sarcosine (w/v), pH 9.5 and incubated overnight at 50 °C and 60 rpm, and washed three times for 15 minutes in 50mM EDTA, pH 8.0 at room temperature and 60 rpm. For chromosome separation, the plugs were transferred to a 1% Certified Megabase agarose (Bio-Rad) gel in 0.5X TBE buffer (Fisher BioReagents™, Madrid, Spain) and subjected to electrophoresis in a CHEF-DR® III system (Bio-Rad) with the following settings: switching time 1,200–4,800 s, 1.6 V/cm, 120 angle, 700 ml/min flow, 8 °C and 255 hours. Gels were then stained for 30 minutes in GelRed® solution (3X final concentration in water; (Biotium, Fremont, California USA) and chromosomal bands were visualized on a GelDoc Go imaging system (Bio-Rad).

Southern blotting was performed as previously described^81^. After the denaturing treatment, chromosomes were transferred from the CHEF gel onto a positively charged nylon membrane (Roche, Barcelona, Spain) and hybridized with digoxigenin-labelled DNA probes generated using the DIG DNA Labelling Mix (Roche). Detection was performed using an alkaline phosphatase kit and CDP-Star® chemiluminescent solution (Roche). Images were obtained with a chemiluminescence detection imager (LAS-3000 Fujifilm, Barcelona, Spain).

### Stress tolerance tests

Based on the functional annotation different environmental stress factors were selected and the response of chromosome 12 mutants to these different environmental stress factors was determined. Strains II5, M1, and selected mutants were inoculated on PDA plates from glycerol stock. Afterwards, agar plugs from the border of the colony were placed on regular PDA plates, MMA plates^82^ or MMA supplemented with either 0.8M NaCl, 1.2M glycerol, 10 µg/mL menadione or 40 µg/mL Congo Red. Colony diameter was measured at 3 dpi. Colonies were photographed at 5 dpi. VCG testing was also performed as described previously^82^.

### *Fusarium* transformation

Modified CRISPR-Cas9 vectors were used for transformation based on the plasmid previously described^83^. This plasmid (CRISPR/Cas9-FoU6-FoNLSx2) contains an additional H2B NLS and an optimized gRNA scaffold^84^. Cloning of the target sequence into the vector and generation of the donor template were performed as previously described^83^. TR4 reference strain II5 and II5 ΔAC12 6.4 were transformed as described^83^ with modifications. Spores of II5 and II5 ΔAC12 6.4 were produced as described below. Subsequently, 500 µL of spore suspension was used to inoculate 50 mL of Fries medium and left to grow in the dark at 135 rpm 20 °C for 1 day. Newly formed mycelium was blended using an IKA® Ultra Turrax® Tube Drive P control (IKA, Staufen, Germany) for 10 seconds at 6000 rpm. All the blended mycelium was used to inoculate 200 mL of fresh Fries medium and incubated overnight in the dark at 135 rpm 20 °C. Overnight formed mycelium was filtered through a layer of nylon mesh (20 micron) and washed three times with 50 mL of KC solution (0.6 M KCl, 65 mM CaCl_2_). Washed mycelium was collected and treated with 10 mL of the filter sterilized protoplasting enzyme mix consisting of 25 mg/ml Driselase from *Basidiomycetes* sp. (Sigma-Aldrich, St. Louis, Missouri USA), 5 mg/ml lysing enzymee from *Trichoderma harzianum* (Sigma-Aldrich, St. Louis, Missouri USA) and 100 μg/ml chitinase from *Trichoderma viride* (Sigma-Aldrich, St. Louis, Missouri USA) in KC. The enzyme mix was incubated at 28 °C with slow rocking movement for two hours. Protoplasts were seperated from the mycelium by filtering through a layer of nylon mesh. Protoplasts were centrifuged at 1800g for 10 minutes at 4 °C and washed twice using ice cold STC (1.2 M Sorbitol, 10 mM Tris-HCL pH 8, 20 mM CaCl_2_). After washing protoplasts were resuspended in 100 µL of STC per reaction. Protoplasts were then mixed with approximately 15 μg of Cas9 plasmid and 15 μg of donor template and 100 µL of 60% PEG solution (Polyethylene glycol 4000, dissolved in STC). Two different gRNA plasmids were combined for knockout of 1g02190. The mixture was left at room temperature for 20 minutes. Protoplasts were washed one last time with 1 mL of STC. Protoplasts were plated by mixing with cooled RM (34.2% sucrose, 0.1% yeast extract, 0.8% agar). After solidifying, a second layer of YG medium (2% glucose, 0.5% yeast extract, 1.5% agar) supplemented with 50 μg/ml nourseothricin was poured over the first agar layer. Resistant colonies were put on separate plates between 3-5 days and used for further genotyping by PCR and amplicon sequencing. Complementation of 12g00300 was performed with the same procedure without Cas9 for ectopic integration.

A II5:Δ*SIX13* mutant containing an hygromycin resistance cassette was generated by a protocol using Cas9 ribonucleoprotein gene editing as described^85^ with modifications. Specifically, the NLS of H2B was cloned from TR4 strain II5 and used to replace the SV40 NLS in the Cas9 plasmid pET-28b-Cas9-His^86^. A hygromycin cassette donor template with short 60 bp homologous overlap was used. Used primers are listed in Table S1. Coding sequence and amino acid sequences of the effectors are listed in Table S2.

### Plant growth conditions and infection assays

Cavendish ‘Grand Naine’ plants were grown in the greenhouse at a day/night temperature of 25/23°C and a day length of 16 hours with 80% relative humidity.

Infection assays of Cavendish ‘Grand Naine’ banana plants were performed as described previously^42^. In brief, strains II5, M1, and selected mutants were inoculated on PDA plates. Agar plugs from PDA plates were used to inoculate Erlenmeyer flasks containing mung bean broth. Flasks were cultivated in a rotary shaker at 150 rpm 25 °C for 5 days. Spores were collected by filtering the culture through two layers of sterile cheesecloth. Spores were subsequently diluted to 1×10^6^ spores per mL. Roots of the plants were injured prior to inoculation. Lastly, 200 mL of spore solution was poured directly on the soil for each plant. Corm necrosis was quantified at 8-10 weeks post inoculation using ImageJ.

## Acknowledgements

The authors thank Li-Jun Ma, University of Massachusetts Amherst, for providing the US copy of TR4 strain II5. JD, ACW, CAG, GNT and GHJK were supported by the Bill and Melinda Gates Foundation, grant no.: AG – 4425. Banana research at Wageningen University has been supported by the Dutch Dioraphte Foundation, grant no.: 20 04 04 02. LGG was supported by grant PLEC2021-007777 from the Spanish AEI/10.13039/501100011033 and NextGenerationEU/PRTR.

