## Supplementary figures 1-14 for "Extensive intrachromosomal duplications in a virulence-associated fungal accessory chromosome"

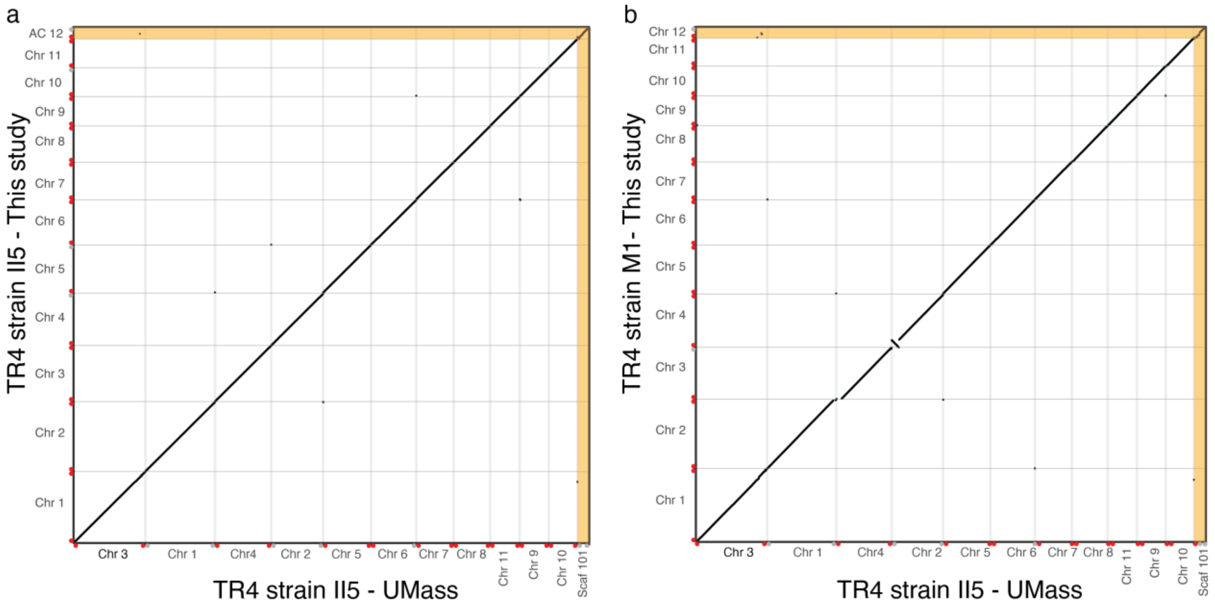


**Fig. S1. – Independently assembled TR4 strains are highly similar.** Whole-genome comparison of TR4 strain II5 (a) and M1 (b) used in this study versus the TR4 strain II5 assembled by Zhang et al. (2024), which had been reported to lack conventional accessory chromosomes. The alignment shows that the assembly of strain II5 used in this study is highly similar to the strain II5 assembled by Zhang et al. (2024). Strain M1 shows a small inversion in chr 4 and a deletion in Chr 3, but furthermore is largely colinear to TR4 strain II5 assembled by Zhang et al. (2024). Importantly, Scaffold101 is homologous to AC12 in our study in both M1 and TR4 II5. Telomeres at chromosomal ends are indicated by red dots, grey dots indicate the absence of detectable telomeric repeats.

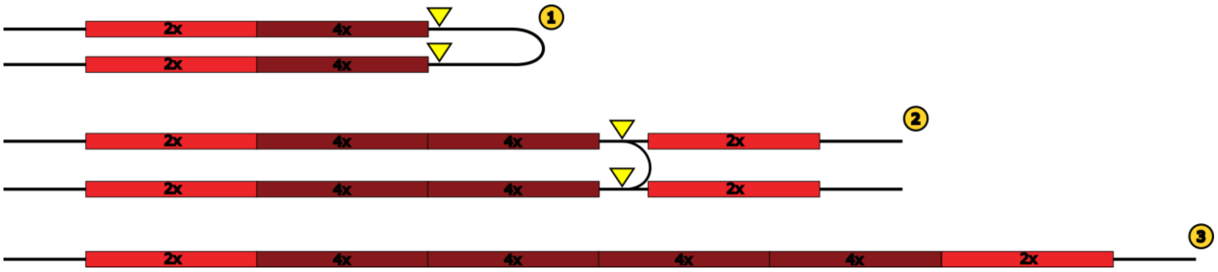


**Fig. S2. – Breakage-fusion-bridge cycles as a potential source of the intrachromosomal duplications of accessory chromosome 12 (AC12).** Schematic representation of a possible mechanism for generation of large intrachromosomal duplications in AC12. Two separate sequential breakage-fusion-bridge events can lead to the displayed order of duplicated and quadruplicated sections of AC12, leading to a final chromosome with triplicate size. Yellow arrows indicate fold-back inversions of sister chromatids after loss of one telomere.

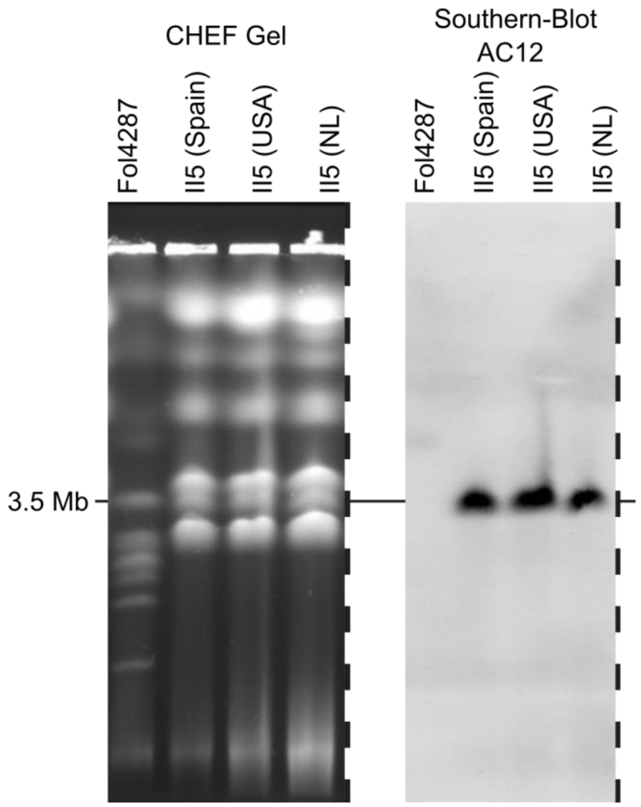


**Fig. S3. – The duplication of accessory chromosome 12 (AC12) in isolate II5 is stable across different laboratories.** CHEF gel of two *Fusarium oxysporum* strains (*Fol*4287 and II5). Three independent copies of the TR4 reference isolate II5 maintained in different laboratories (Spain, USA, and NL) were tested. The same gel was used for a Southern Blot using an AC12 specific probe. Dashes indicate cropping from gel. Full gel depicted in Fig. S14.

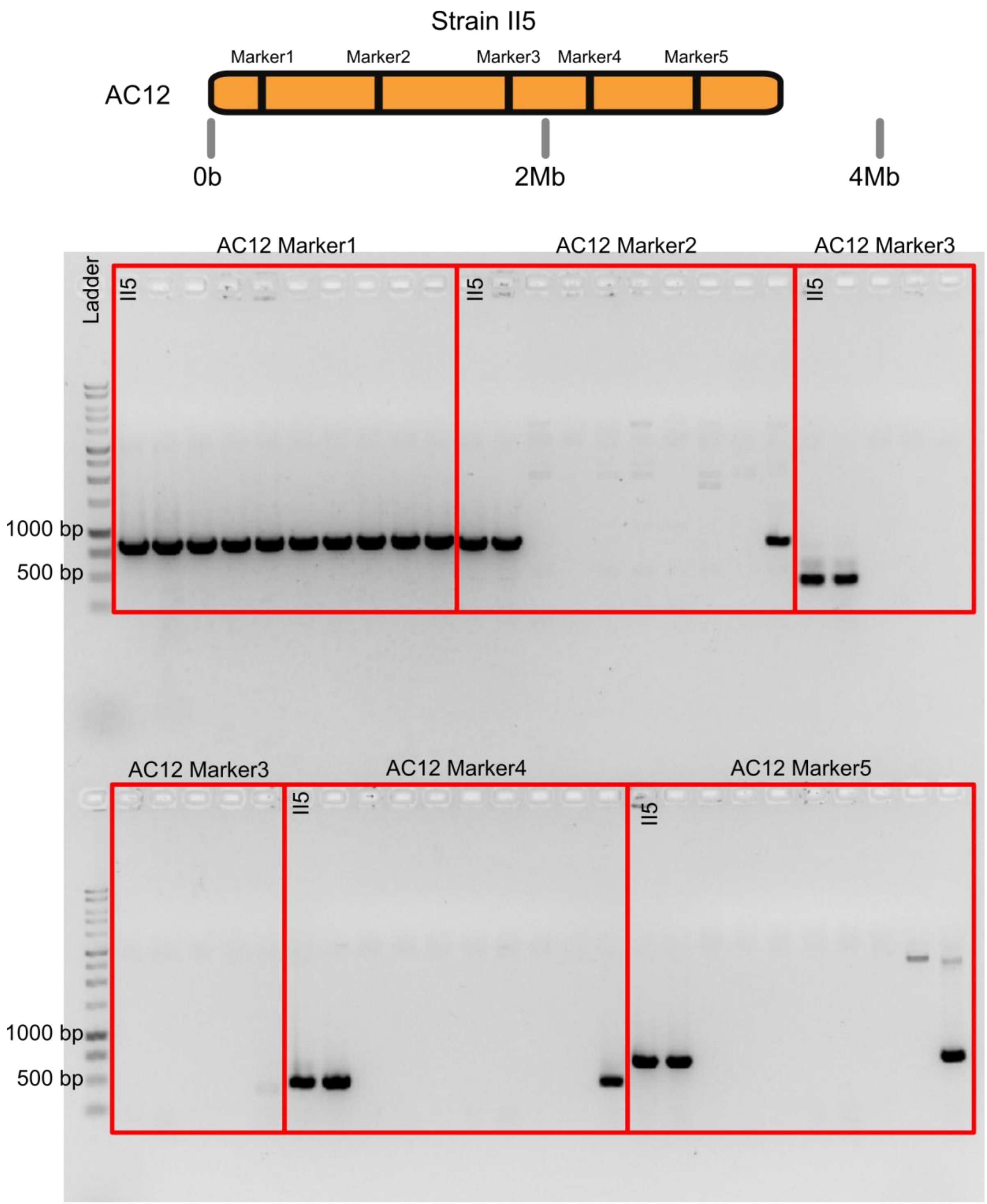


**Fig. S4. - PCR indicates loss of accessory chromosome 12 (AC12) after benomyl treatment.** Schematic representation of primer locations on AC12 (black bands) and gel electrophoresis after PCR on strain II5 and putative AC12 mutants using five different AC12 PCR markers. Seven colonies did not give bands for markers 2-5. Note, marker 1 can also bind to the accessory region on chromosome 1, therefore giving a band for all colonies.

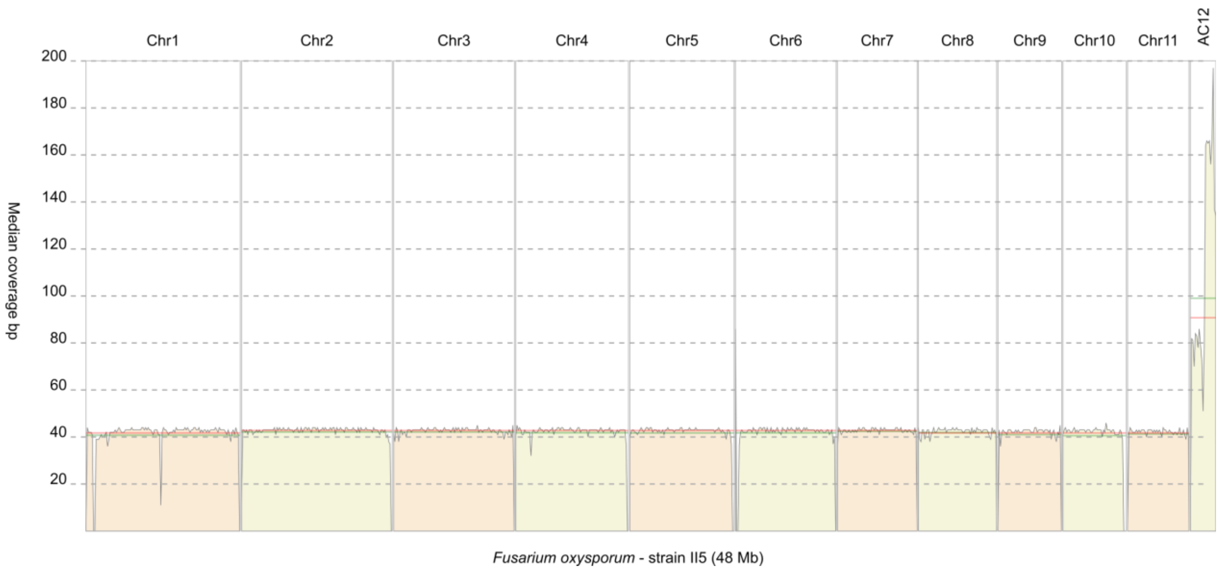


**Fig. S5. - Mapping of short reads to the II5 reference genome assembly.** Mapping of Illumina reads from strain II5 to the II5 reference genome assembly shows higher coverage at accessory chromosome 12 (AC12).

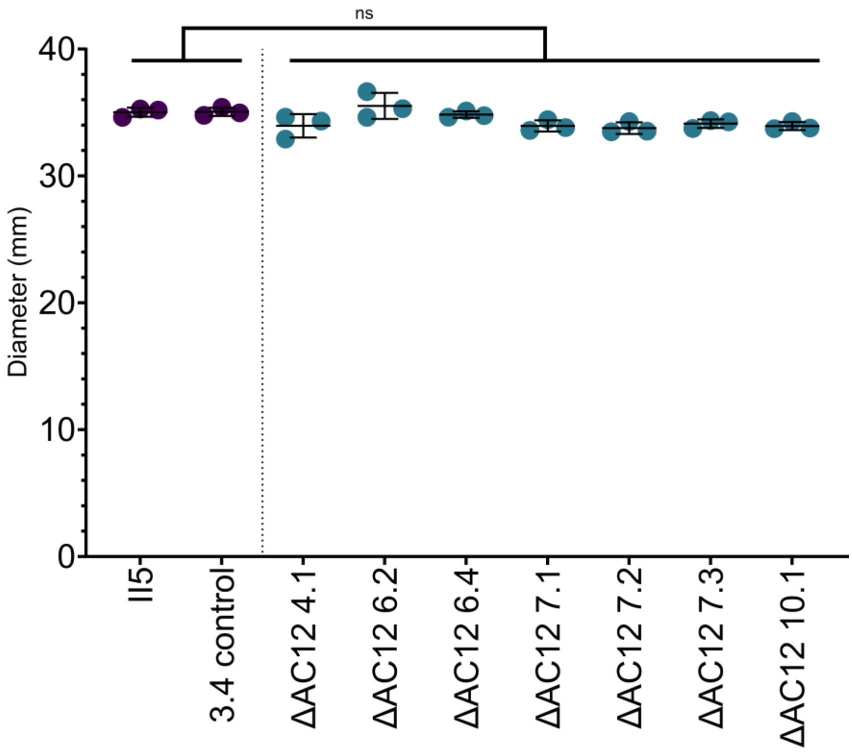


**Fig. S6. – Loss of accessory chromosome 12 (AC12) does not affect vegetative growth on PDA.** Colony diameter (mm) of PDA plates inoculated with TR4 strain II5, benomyl treated control 3.4, and seven independent AC12 mutants. Growth was quantified at 6 dpi. Letters indicate significant differences between treatments (Tukey-Kramer test; $P<0.05$). Data are shown as mean ±SD (n=3).

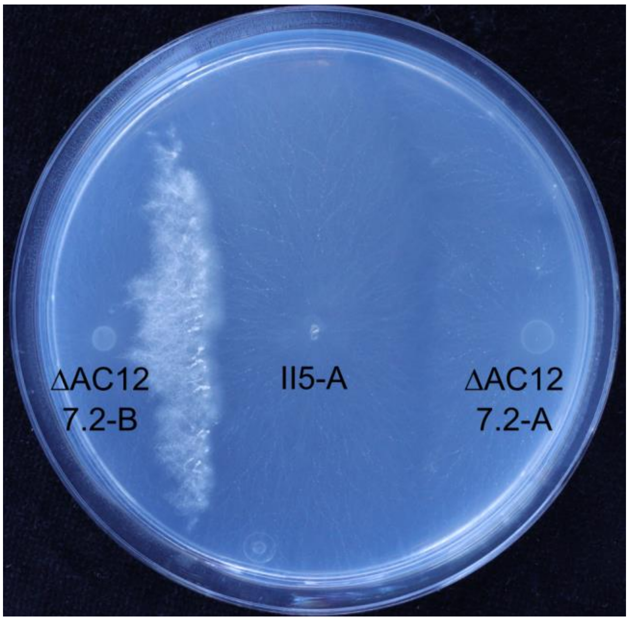


**Fig. S7.** - **Accessory chromosome 12 (AC12) mutant is vegetatively compatible with WT strain II5.** MMA plate inoculated with *nit* mutant of II5 and II5ΔAC12 7.2. Dense hyphal growth at the colony contact points indicates vegetative compatibility.

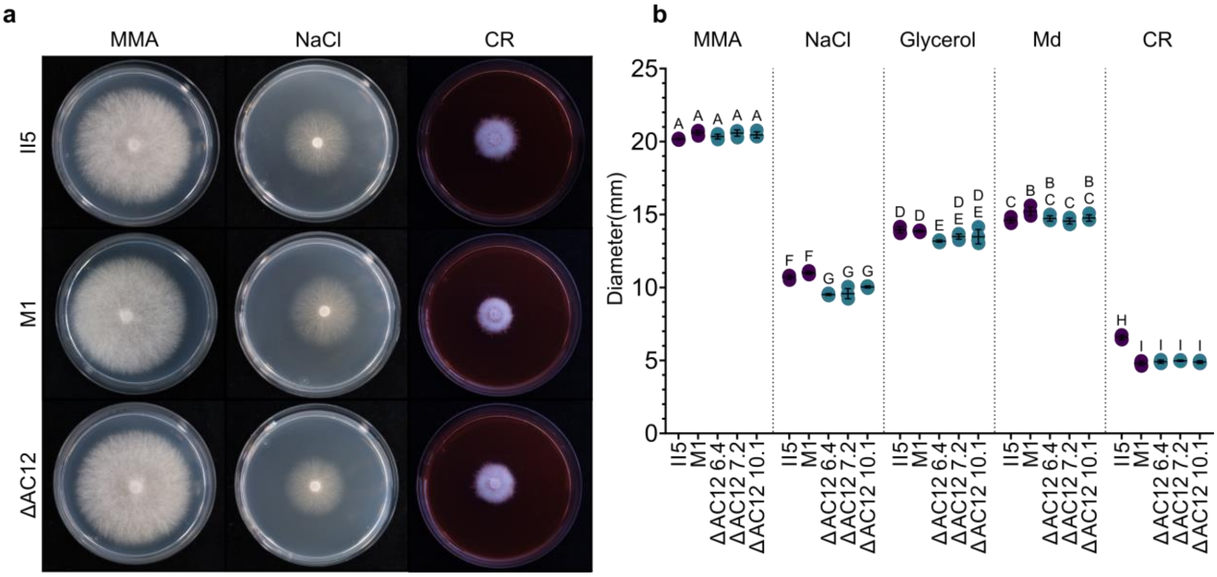


**Fig. S8. - Accessory chromosome 12 (AC12) plays a role in growth under different stress factors. a**) MMA plates alone or supplemented with 0.8 M NaCl or 40 µg/mL Congo Red (CR) were inoculated with TR4 strains II5, M1, or II5ΔAC12 6.4 and photographed at 5 dpi. **b**) Colony diameter (mm) of MMA plates alone or supplemented with 0.8 M NaCl, 1.2 M glycerol, 10 µg/mL menadione (Md) or 40 µg/mL Congo Red (CR) inoculated with TR4 strains II5, M1 or three independent AC12 mutants. Colony growth was quantified at 3 dpi. Different letters indicate significant differences between treatments (Tukey-Kramer test; $P<0.05$). Data are shown as mean ±SD (n=4).

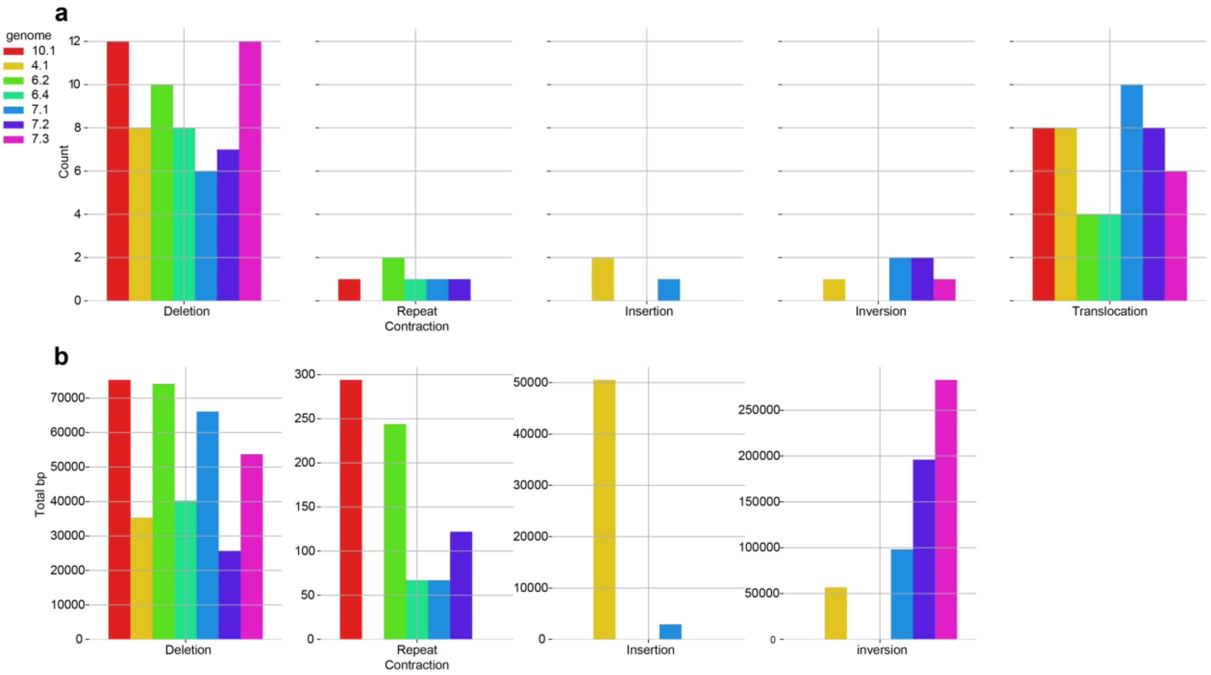


**Fig. S9. - Structural variants in the accessory chromosome 12 (AC12) deletion mutants. a**) The number of structural variants per variant type is shown for all seven independent AC12 deletion mutants. **b**) Sizes (in bp) of the identified structural variants for all seven independent AC12 deletion mutants. Translocation sizes are not reported because the calculation is challenged by the fragmented short read assemblies.

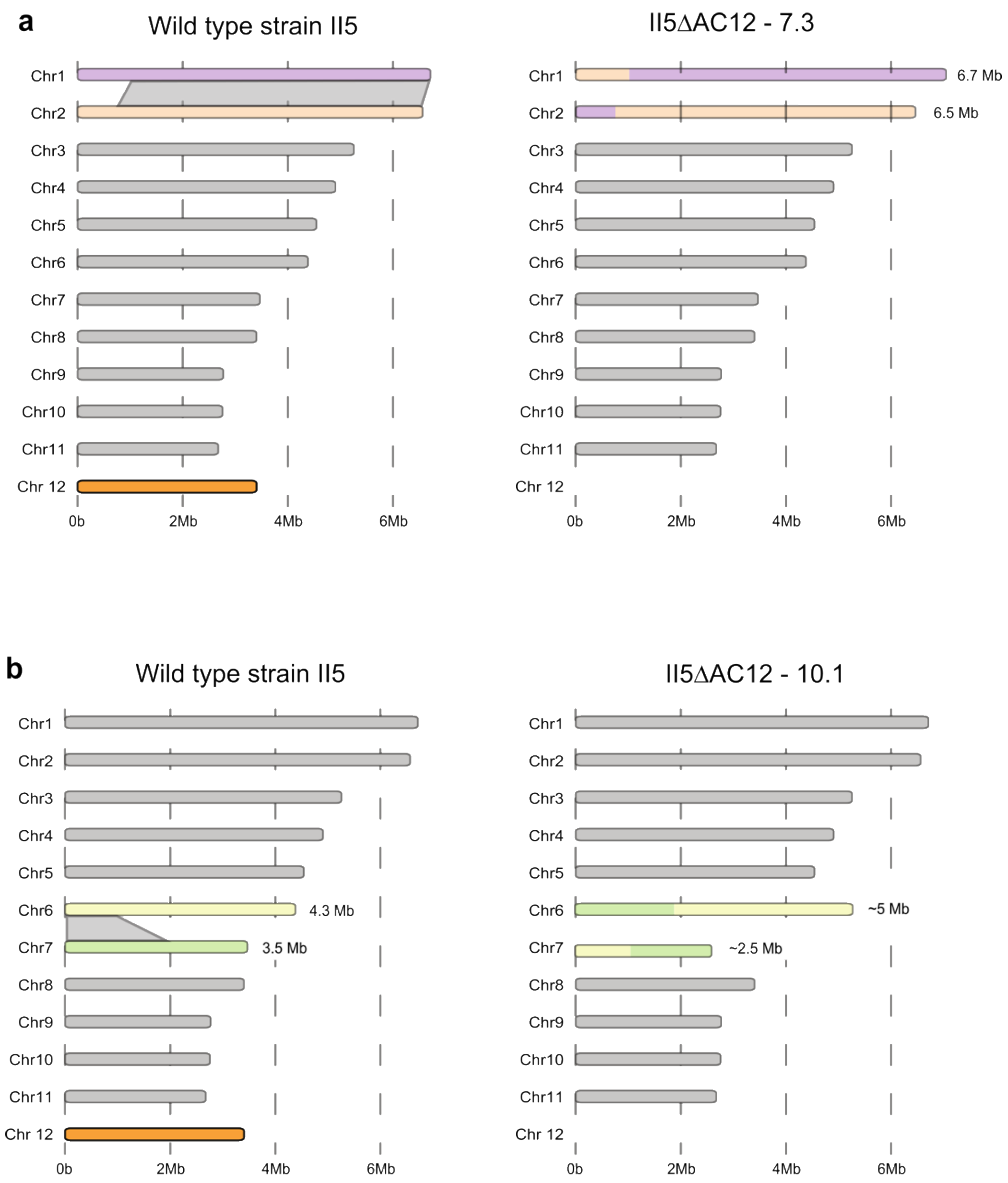


**Fig. S10. - Interchromosomal rearrangements of core chromosomes following benomyl treatment and accessory chromosome (AC12) loss. a**) Schematic representation of the interchromosomal rearrangements that occurred in II5ΔAC12 - 7.3. Translocations between chromosomes 1 and 2 are highlighted by ribbons. **b**) Schematic representation of the interchromosomal rearrangements that occurred in II5ΔAC12 - 10.1. Translocations between chromosomes 6 and 7 are highlighted by ribbons.

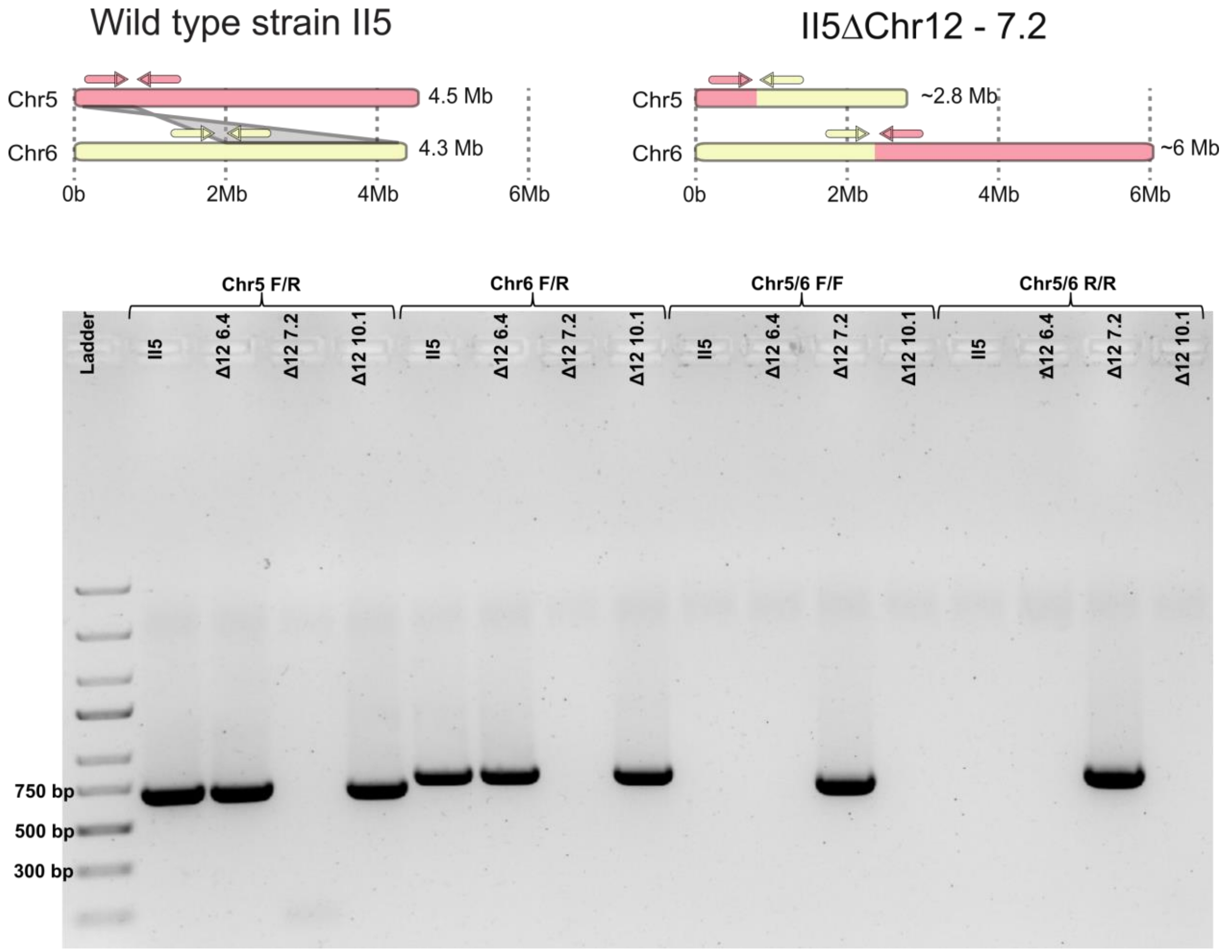


**Fig. S11. Core chromosomal rearrangement in II5ΔAC12 - 7.2 are confirmed through PCR.** Schematic representation of primer locations on chromosomes 5 and 6 in II5 and II5ΔAC12 - 7.2. Gel electrophoresis after PCR on II5 and three independent chromosome 12 mutants using chromosome 5 (F/R) and 6 (F/R) specific primers and the combinations thereof.

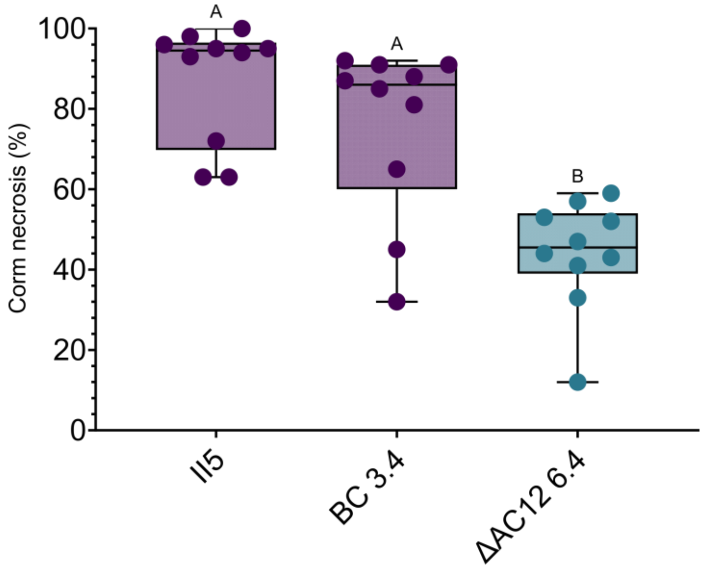


**Fig. S12. - Benomyl treatment does not affect virulence.** Percentage of corm necrosis of Cavendish 'Grand Naine' plants inoculated with strain II5, benomyl treated control 3.4 and II5ΔAC12 6.4. Corm necrosis was quantified using ImageJ (n=10). Letters indicate significant differences between treatments (Tukey-Kramer test; P<0.05).

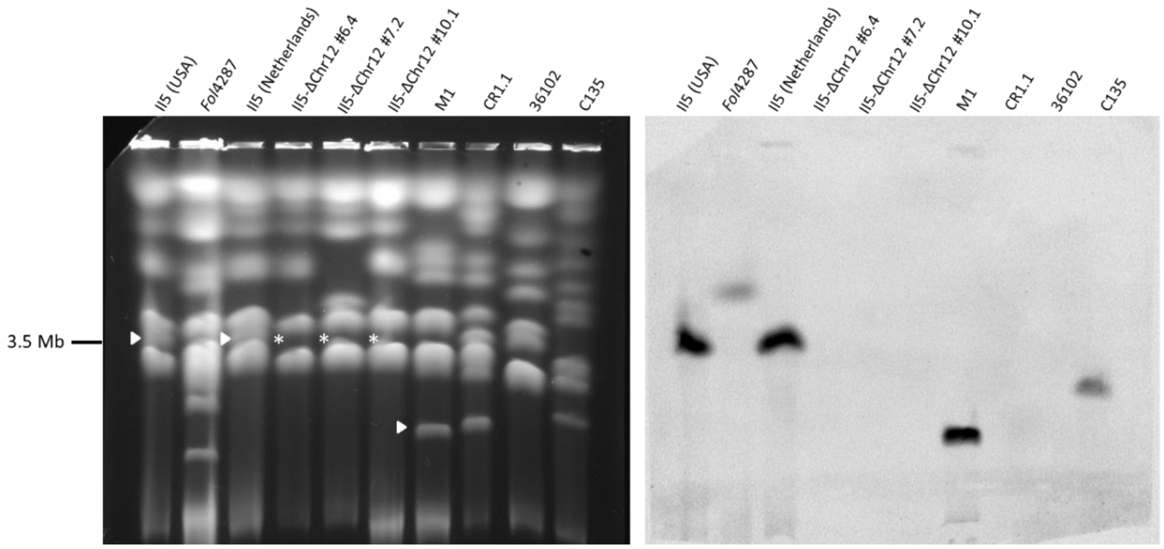


**Fig. S13. -** Full uncropped CHEF gel and Southern blot used in Fig. 1 and 2.

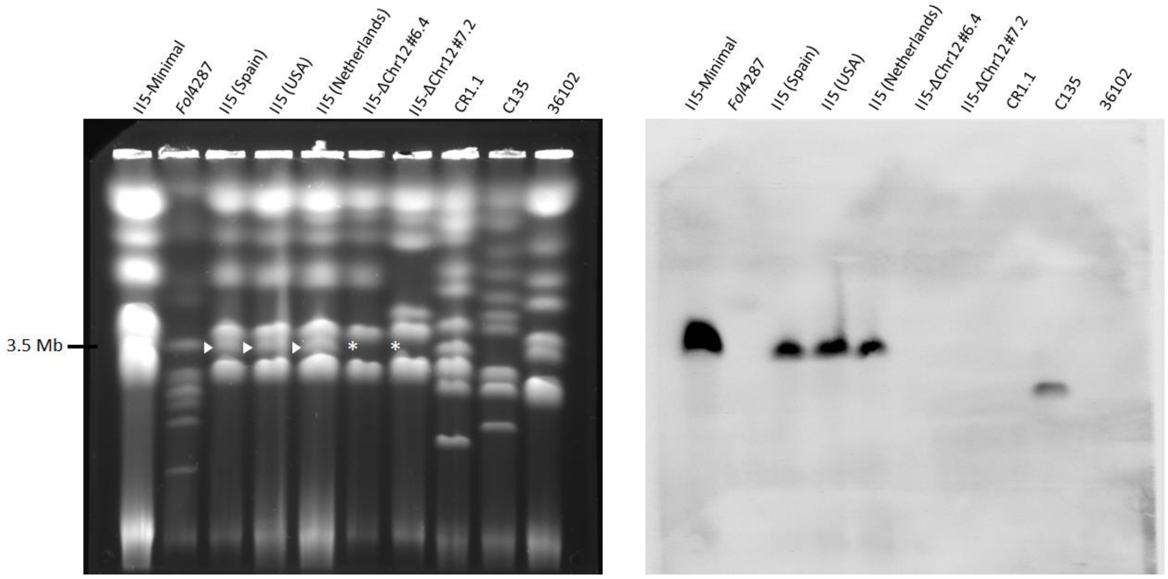


**Fig. S14. -** Full uncropped CHEF gel and Southern blot used in Fig. S3. II5-Minimal concerns an isolate for an unrelated study.
